# Spreading pre-saccadic attentional resources without trade-off

**DOI:** 10.1101/187518

**Authors:** Michael Puntiroli, Heiner Deubel, Martin Szinte

## Abstract

When preparing a saccade, attentional resources are focused at the saccade target and its immediate vicinity. Here we show that this does not hold true when saccades are prepared towards a recently extinguished target. We obtained detailed maps of orientation sensitivity when participants prepared a saccade toward a target that either remained on the screen or disappeared before the eyes moved. We found that attention was mainly focused at the immediate surround of the visible target and increasingly spread to more peripheral locations as a function of the delay between the target’s disappearance and the saccade. Interestingly, this spread was accompanied by an overall increase in sensitivity, speaking against a dilution of limited resources over a larger spatial area. We hypothesize that these results reflect the behavioral consequences of the spatio-temporal dynamics of visual receptive fields in the presence and in the absence a structured visual cue.

## Introduction

To efficiently make sense of our rich visual environment, the visual system evolved and gained the ability to selectively process the most salient information (Itti & Koch, 2001). This selection is, however, limited by the architecture of the visual system itself (Anton-Erxleben & Carrasco, 2013). To compensate for the low visual resolution in peripheral vision, selection can either be achieved by shifting high resolution central vision to peripheral objects of interest by means of saccades (overt attention), or by shifting spatial attention while keeping the eyes steady (covert attention). Saccades are preceded by a shift of attention toward the saccade target (e.g. Armstrong & Moore, 2007; Deubel & Schneider, 1996; Moore & Fallah, 2004) suggesting that both overt and covert attention rely on similar processes (Awh, Armstrong, & Moore, 2006; Rizzolatti, Riggio, & Sheliga, 1994). Indeed, both cases result in the deployment of attention resources, leading to spatially localized gains in reaction time (e.g. Posner, 1980; Remington, 1980), in visual sensitivity (e.g. Bashinski & Bacharach, 1980; Deubel & Schneider, 1996) and in neural activity (e.g. Moran & Desimone, 1985; Wurtz & Mohler, 1976).

Moreover, it has been shown that neurons are particularly sensitive to stimuli presented in the direction of an attended target (saccade target or cue) located outside their classical receptive fields (Anton-Erxleben, Stephan, & Treue, 2009; Connor, Preddie, Gallant, & Van Essen, 1997; Moran & Desimone, 1985; Neupane, Guitton, & Pack, 2016; Niebergall, Khayat, Treue, & Martinez-Trujillo, 2011; Tolias et al., 2001; Womelsdorf, Anton-Erxleben, & Treue, 2008; Womelsdorf, Anton-Erxleben, Pieper, & Treue, 2006; Zirnsak, Steinmetz, Noudoost, Xu, & Moore, 2014). Such attentional modulation of the visual neurons’ spatial tuning is accompanied by changes in their receptive fields size and position, as observed in animal electrophysiology (Anton-Erxleben et al., 2009; Womelsdorf et al., 2006; 2008) and in humans, using functional imaging methods (Herrmann, Montaser-Kouhsari, Carrasco, & Heeger, 2010; Kay, Weiner, & Grill-Spector, 2015; Klein, Harvey, & Dumoulin, 2014; Sprague & Serences, 2013). The allocation of attention can then be modeled as a gaussian field of variable width, centered on the attended cue location, modulating the spatial tuning of visual cells’ receptive fields located nearby (Reynolds & Heeger, 2009; Womelsdorf et al., 2008). Furthermore, the width of this gaussian field is thought to be essential, as it determines the type of attentional benefits observed (contrast gain vs. response gain) in visual sensitivity (Herrmann et al., 2010) and in neural activity (Carandini & Heeger, 2012; Reynolds & Heeger, 2009).

Attention then certainly modulates visual cells’ receptive fields, but what are the consequences at the behavioral level of such modulation? If it is the presence of an attended target that drives the modulation of the visual cell’s receptive field tuning, what happens after its disappearance? To answer these questions we will first have to clarify how the spatial modulation of attention can be assessed behaviorally. Different authors have assessed the spatial spread of attentional benefits through measures of reaction time or visual sensitivity change at multiple location surrounding a cue. When testing covert attention during fixation attentional benefits were found at positions extending over the whole visual field tested (Tse, Sheinberg, & Logothetis, 2003), or at positions limited to one visual hemifield (Hughes & Zimba, 1985), to a visual quadrant (Hughes & Zimba, 1987) or to a few degrees surrounding the cue (Henderson & Macquistan, 1993; Shulman, Remington, & McLean, 1979; Shulman, Wilson, & Sheehy, 1985). These huge variations seem to primarily reflect methodological discrepancies, such as the use of exogenous or endogenous cues, and also variations in the task difficulty (Intriligator & Cavanagh, 2001). Indeed, by using a structured visual field (Eriksen & Yeh, 1985; Taylor, Chan, Bennett, & Pratt, 2015), by increasing the difficulty of the tasks by masking the targets (Deubel & Schneider, 1996; Baldauf:2006hn; Doré-Mazars, Pouget, & Beauvillain, 2004; Handy, Kingstone, & Mangun, 1996; Henderson, 1991; Kowler, Anderson, Dosher, & Blaser, 1995), or by controlling for visual eccentricity effects (Koenig-Robert & VanRullen, 2011d), it was shown that the attentional spread was narrowly concentrated, with benefits limited to a few degrees of visual angle surrounding a covertly attended cue or a saccade target.

Using a new visual sensitivity mapping paradigm inspired from the above work, we evaluated the spatial extent of attentional benefits before the execution of a saccade toward a cue. In particular, we evaluated the spatiotemporal dynamics of attention following the disappearance of the cue. We found that highest discrimination sensitivity was concentrated at the cue location and its immediate surrounds when the cue was visible. However, after the cue’s disappearance, benefits spread to more peripheral locations. This spread was accompanied by an overall increase in sensitivity and cannot be explained by a loss in spatial localization of the memorized location. As this spread of attention was made without any evidence of a trade-off of attentional resources (meaning it was not accomplished by spreading limited resources over a larger spatial area), our effects revealed a yet unknown consequence of attention field modulation. In particular, they suggest that visual neuron’s receptive fields shift toward the visible cue, but when the cue is no longer visible, return to their initial coordinates, accompanied by a spatial spread of attention benefits without any attentional trade-off.

## Results

Our goal was to determine the spatial distribution of attentional benefits when participants prepared a saccade toward a cue that either remained on the screen, or had recently disappeared. To this end, we probed attention by presenting a discrimination target at one of various locations surrounding the cue (Figure 1). Through the use of a threshold task, we kept discrimination performance homogeneously high across space despite the fact that discrimination targets appeared at several eccentricities from the fixation target (see experimental procedure and Figure 1D). Then, to study the spatio-temporal dynamics of attention following the disappearance of the cue, we systematically varied the delay between the initiation of the saccade and the disappearance of the cue.

**Fig. 1.**
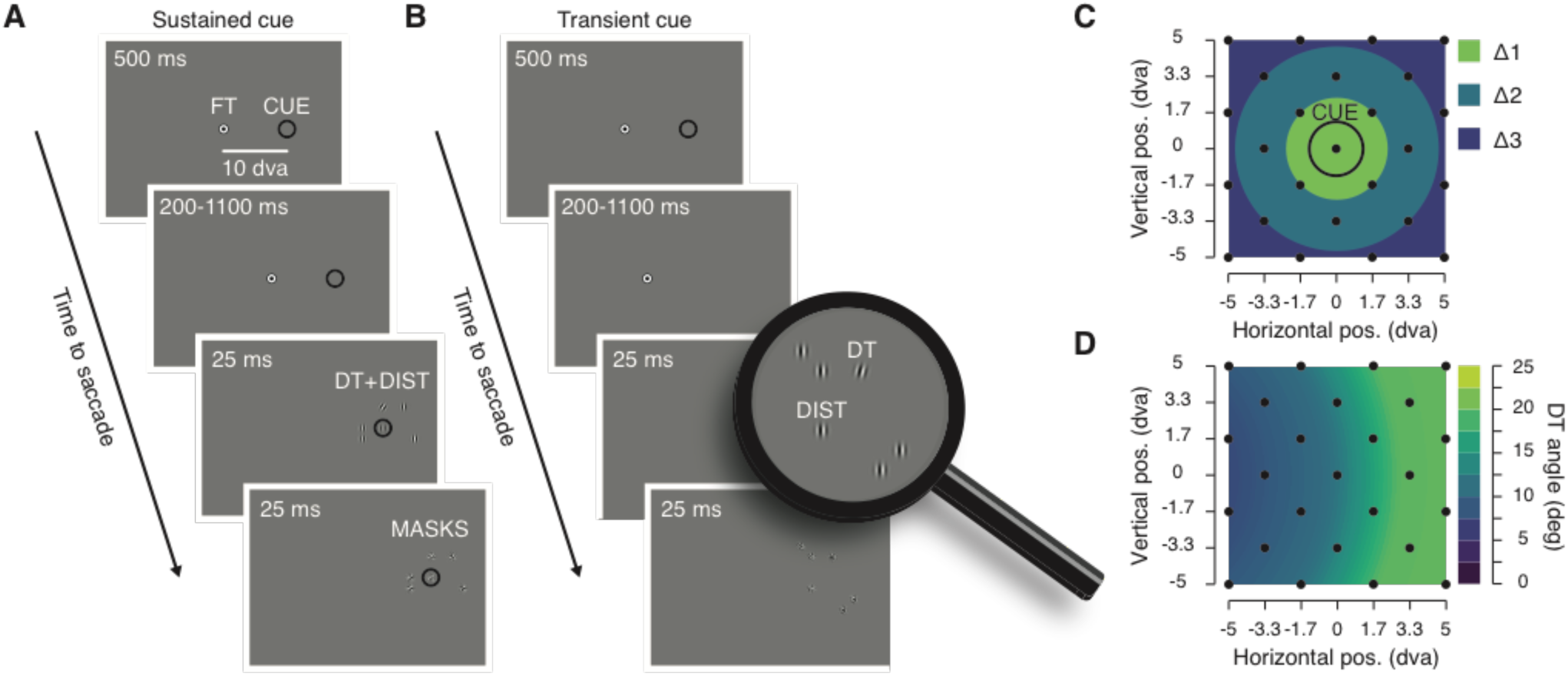
Experimental procedure. **A**. Sustained cue condition. Participants prepared a saccade from the fixation target (FT) to a visual cue (CUE) presented continuously on the screen throughout the trial. Participants were instructed to saccade toward the center of the cue at the offset of the FT, which occurred between 700 and 1600 ms after the cue onset. Just before the saccade, a discrimination target was shown (DT, 25 ms clockwise or counter-clockwise tilted Gabor) together with 5 distractors (DIST, vertical Gabors) and followed by 6 overlaying masks (MASKS, 25 ms noise patches). **B.** Transient cue condition. Participant prepared a saccade from the FT to a CUE presented transiently (500 ms). Participants saccade at the offset of the FT which occurred between 200 and 1100 ms after the cue offset. **C.** On each trial the position of the DT and of the distractors were randomly picked between 25 possible positions (black dots), homogeneously covering a 10° by 10° map centered on the CUE. **D.** Before the main saccade task, we determined at different eccentricities from the FT the necessary DT angle leading to a correct discrimination level of 80%. The graph shows averaged DT angle (n=12) interpolated across the different DT eccentricities from the FT (see Method). DT angle is shown via the color scale.

We first verified that the presentation of the discrimination target itself did not systematically influence oculomotor behavior. We did not find any differences with respect to saccade latency when comparing trials with and without the presentation of a discrimination target (4% of trials were without discrimination target, present: 201.77 ± 2.84 ms vs. absent: 199.96 ± 7.12 ms, *p*= 0.1726) and only a slight change in saccade amplitude (present: 9.67 ± 0.17° vs. absent: 9.77 ± 0.40°, *p* < 0.0366), therefore validating our procedure. We next obtained maps of visual sensitivity, reflecting participants’ ability to correctly report the orientation of the discrimination target presented at different distances from the cue. Figure 2 shows sensitivity maps obtained across participant by presenting discrimination targets just before the saccade at 25 different positions (see Figure 1C), for trials in which the cue remained on the screen (Figure 2A) and trials in which it disappeared between 200 ms and 1100 ms before the saccade (Figure 2B-D). Despite the limited amount of trials obtained per participants for each of the tested positions (40.76 ± 1.26 trials), the maps make it possible to appreciate the effects of the cue on the allocation of attention. Indeed, they show that attentional benefits were more pronounced toward the immediate contour of the cue and subsequently spread, more and more, as the delay between the saccade onset and the cue offset increased. These effects were systematically analyzed by combining the 25 tested positions into 3 groups of discrimination target distances from the cue (see Δ1, Δ2 and Δ3 in Figure 1C). Within the trials in which the cue remained on the screen (Figure 3A), performance was best for discrimination targets presented within ~2.4°surrounding the cue (Δ1: d#x2019; = 1.55 ± 0.18 vs. Δ2: d#x2019; = 1.03 ± 0.15, *p* < 0.0001; Δ1 vs. Δ3: d’ = 0.92 ± 0.11, *p* < 0.0001), with sensitivity at the immediate surround of the cue being approximately 63% higher compared to discrimination targets shown at further distances (Δ1/Δ2: 159.54 ± 14.65%, Δ1/Δ3: 170.97 ± 13.14%). These results are in line with previous evidence showing that the pre-saccadic shift of attention is limited toward the closest positions surrounding a saccade target (Deubel & Schneider, 1996).

**Fig. 2.**
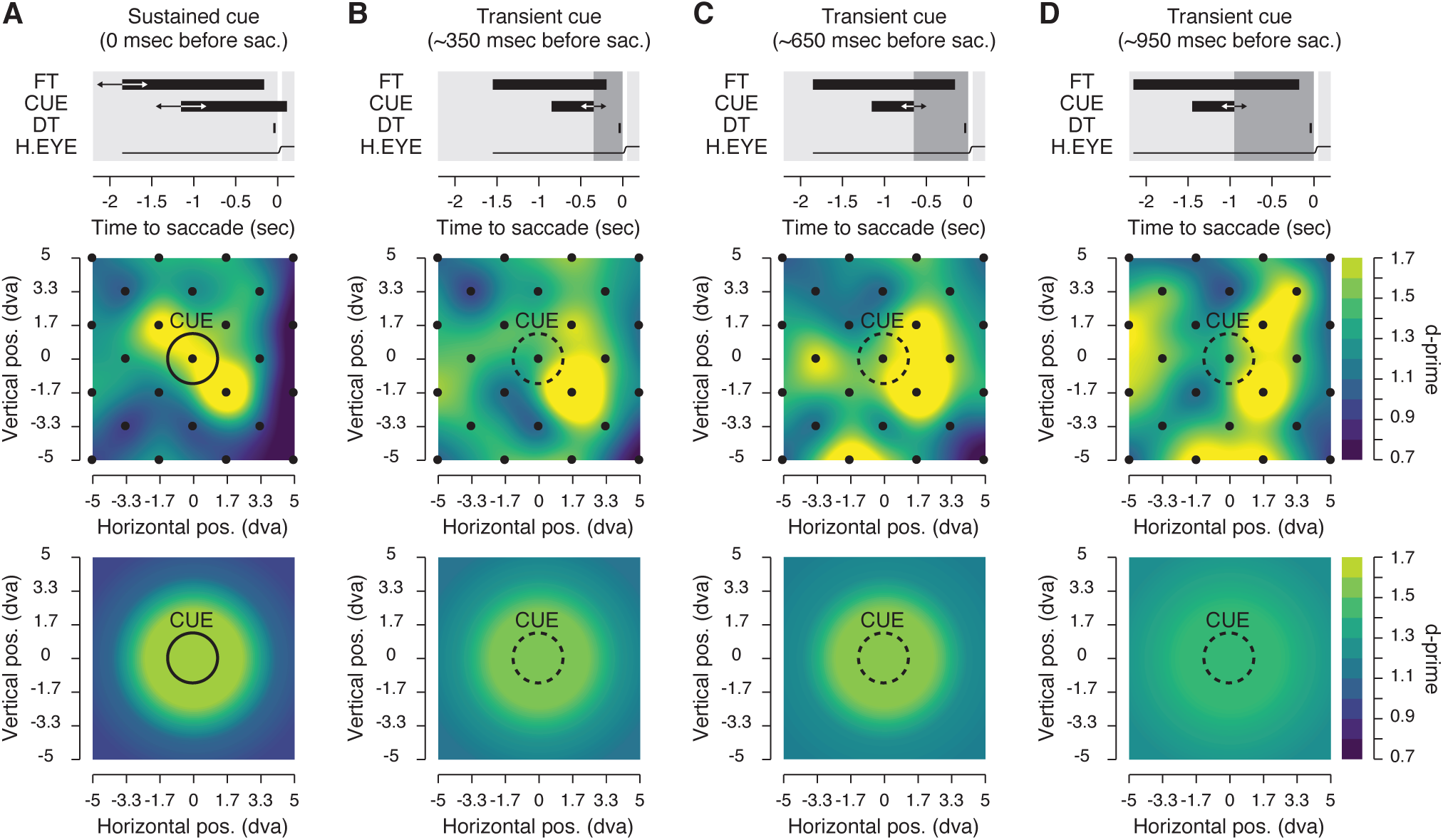
Pre-saccadic sensitivity maps. Each graphs shows average sensitivity across all participants (see Method) gathered either at 25 positions individually (middle row) or grouped in 3 distances (Δ1, Δ2 and Δ3, bottom row) surrounding the saccade cue (CUE). The top row of each panel describe the time course relative to the saccade onset of the fixation target (FT), the cue (CUE), the discrimination target (DT) and the horizontal eye position (H. EYE). Data are shown for the sustained cue condition (**A**) and the transient cue conditions (**B-D**). The transient cue condition is binned in three equal groups of trials where the cue offset preceded the saccade onset by approximately 350 (B), 650 (C) or 950 ms (D). Averaged sensitivity (d’) is shown via the color scale.

**Fig. 3.**
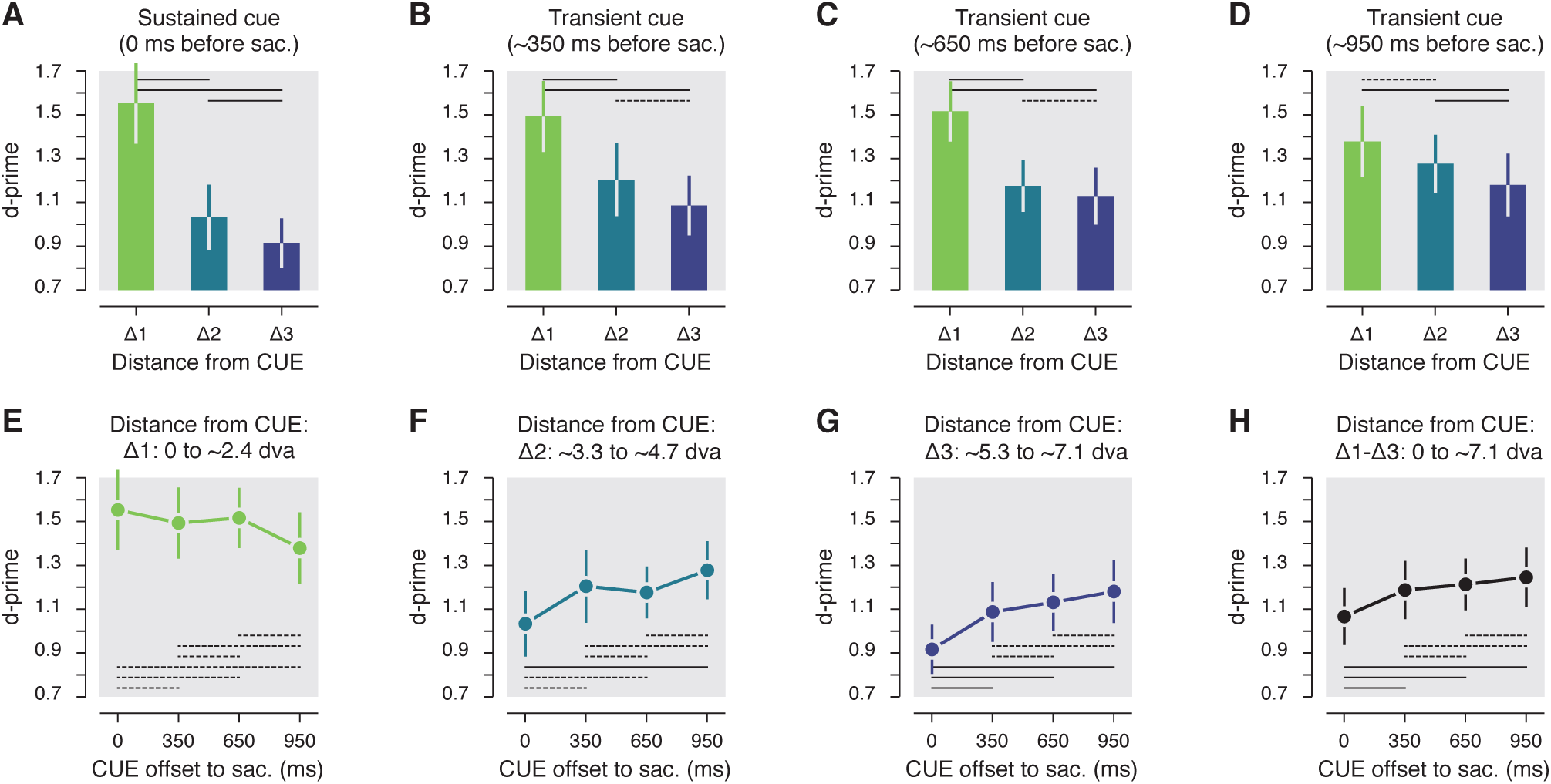
Statistics. **A-D.** Pre-saccadic sensitivity as a function of the distance from the cue center (Δ1-Δ3). Data are shown for the sustained cue (A) and the transient cue conditions (B-D). The transient cue condition is binned in three equal groups of trials where the cue offset precede the saccade by approximately 350 ms (B), 650 ms (C) or 950 ms (D). **E-H.** Pre-saccadic sensitivity as a function of the the duration between the cue offset and and saccade onset. Data are shown separately for three main distance of the DT from the CUE center (E-G) or for all trials irrespective of their distance from the cue (H). Note that overall sensitivity results (H) is not directly the mean of the sensitivity observed at the different distances from the cue (E-G) as there was not the same amount of trials played at each analyzed distance from the cue. Error bars show SEM, dashed and full lines represent non-significant (*p* > 0.05) and significant (*p* < 0.05) comparisons, respectively.

Next, we found a similar pattern of results when, rather than remaining continuously on the screen, the cue was extinguished approximately 350 ms before the saccade (Figure 3B-D). Within these trials we observed a similar distribution of pre-saccadic attention with best performance limited within the immediate contour of the cue (Δ1: d’ = 1.49 ± 0.16 vs. Δ2: d’ = 1.21 ± 0.17, *p* < 0.0406; Δ1 vs. Δ3: d’ = 1.09 ± 0.14, *p* < 0.0004) with an increase of sensitivity of approximately 42 and 46% when compared with performance at the two further distances respectively (Δ1/Δ2: 142.42 ± 22.43%, Δ1/Δ3: 146.49 ± 19.17%). These results seem to indicate that the offset of the cue had no significant influence on the allocation of attention before a saccade, at least within the first 350 ms following its offset. Next, when the cue had disappeared from the screen approximately 650 ms before saccades start (Figure 3C), we found that targets were better discriminated if they were presented within the first distance from the cue (Δ1: d’ = 1.52 ± 0.14 vs. Δ2: d’ = 1.18 ± 0.12, *p* < 0.0008; Δ1 vs. Δ3: d’ = 1.13 ± 0.13, *p* <0.0002), with however a reduced spatial cueing effect (Δ1/Δ2: 137.98 ± 17.56%, Δ1/Δ3: 146.56 ± 13.83%). The trend toward an increasing spread of attention became evident when the cue had disappeared approximately 950 ms before the saccade (Figure 3D). For these trials, the sensitivity difference between first and the second distances from the cue was no longer significantly (Δ1: d’ = 1.38 ± 0.16 vs. Δ2: d’ = 1.28 ± 0.13, *p* = 0.3608) and cueing effects between the first and the two other distances were strongly reduced (Δ1/Δ2: 108.03 ± 11.52%). Also, within these trials, performance for discrimination targets presented between ~3.3° and ~4.7° from the cue (Δ2) was now slightly better than for trials in which they were shown even further away (Δ3) from the cue (Δ2 vs. Δ3: d’ = 1.18 ± 0.14, *p* <0.0484); an effect observed for trials in which the cue remained onscreen (*p* <0.0186) but not for trials in which the cued disappeared less than approximately 950 ms before the saccade (all *ps* > 0.4964).

The results above suggest that attention was, in general, drawn toward the cue and its close proximity, at least when it remained on the screen, or when it disappeared less than 950 ms before the saccade. To capture the time course of the attentional spread, we compared sensitivity gathered at each of discrimination target-to-cue distances (Δ1-Δ3) in function of the delay between the saccade onset and the cue offset. We considered trials in which the cue remained on the screen as the shortest delay (t0, “t” corresponding to the cue offset to the saccade onset time). We found that sensitivity for discrimination targets shown at the cue’s immediate contour (Δ1) remained at a very similar level over the tested delays (0.8324 > *ps* > 0.1102, see dashed lines in Figure 3E). On the other hand, sensitivity for discrimination targets shown at positions more than 2.4° from the cue (Δ2 and Δ3) gradually improved as the time from cue offset increased (Figure 3F-H). In particular, we found a significant improvement of sensitivity for discrimination targets shown between ~3.3° and ~4.7° from the cue (Δ2, Figure 3F) when comparing trials where the cue remained on the screen to trials when the cue offset preceded the saccade by approximately 950 ms (t0: d’ = 1.03 ± 0.15 vs. t950: d’ = 1.28 ± 0.13, *p* < 0.0001), but not when the delay was shorter (i.e. t350 and t650: 0.1256 > *ps* > 0.1076). Such a spread of attention for discrimination target shown at the greatest tested distance from the cue (Δ3, Figure 3G) was visible even earlier in time (i.e. for shorter cue offset to the saccade). Significant differences were, in fact, found in trials where the cue disappeared approximately 350 ms (t0: d’ = 0.92 ± 0.11 vs. t350: d’ = 1.09 ± 0.14, *p* <0.0098) and even earlier relative to the saccade onset (i.e. t650 and t950: both *ps* <0.0001). Interestingly, when considering all discrimination target positions together (Figure 3H), we found that sensitivity increased as a function of the delay between the saccade onset and the cue offset. Overall sensitivity (computed as the average across the 25 possible positions of the discrimination target) increased by approximately 27% when comparing trials in which the cue disappeared about a second before the saccade to those in which the cue remained on the screen (t950/t0: 126.67 ± 10.62%). Furthermore, this increase of sensitivity was already significant when comparing trials where the cue remained on the screen with trials where the cue was extinguished approximately 350 ms before the saccade (t0: d’ = 1.07 ± 0.13 vs. t350: d’ = 1.19 ± 0.13, *p* <0.0490) or even earlier (t0 vs. t650: d’ = 1.21 ± 0.12, *p* <0.0022, t0 vs. t950: d’ = 1.25 ± 0.14, *p* <0.0001).

Altogether, we observed that pre-saccadic attention was captured by the cue and maintained within its close proximity even one second after its disappearance. Moreover, the cue’s disappearance was quickly followed by an improvement of sensitivity at distances further away from it. Such benefits then cannot be explained by a tradeoff of attention resources between the closest and furthest distances from the cue. Rather our results suggest the existence a mechanism allowing a spread of attention over space without a significant loss at the closest positions surrounding the cue.

Can these results be explained by the fact that, after a rather long delay, participants lose track of the cue location? Because we investigated allocation of covert attention in a saccade task, our design allowed us to use saccade metrics to test this alternative explanation. First, we looked at whether the spread of attention was accompanied by a comparable spread of saccade endpoints. Figure 4A-D show the normalized saccade landing frequency for the different delays between the cue offset relative and the saccade onset. From these graphs, one can appreciate the absence of any strong difference in the frequency of saccade landing, which would be expected from the spread of attention observed above. The disappearance of the cue, nevertheless, had a clear influence on the saccades. In particular, we found that saccades were more accurate (accuracy assessed as the absolute distance between the saccade offset and the saccade target, t0: 1.24 ± 0.04° vs. t350-950: 1.47 ± 0.05°, *p* <0.0001) and more precise (precision assessed as the standard deviation of the accuracy, t0: 0.68 ± 0.02° vs. t350-950: 0.80 ± 0.02°, *p* <0.0001) when the cue remained on the screen compared to trials where it disappeared from the screen. We attributed the increased spread of saccade endpoints to the offset of the cue itself (Deubel, Wolf, & Hauske, 1982), and the lack of visual feedback just before saccade onset. In fact, contrary to what we found for attention, when the cue disappeared from the screen both saccade accuracy and precision didn’t change as a function of the delay between cue offset and saccade onset (0.5882 > *ps* > 0.1144). Next, using saccade endpoint coordinates we re-encoded the discrimination target positions in order to obtain sensitivity maps relative to the saccade endpoints, rather than relative to the cue position (Figure 4E-H). We hypothesized that if the spread of attention was due to a spread in saccade landing, then, by correcting the discrimination target coordinates for the saccade endpoints, the differences in sensitivity observed for different delays between cue offset and saccade onset should no longer be found, or at least be reduced. This was not the case. Instead, we found a very similar spread of attention as above, with sustained sensitivity over time observed at the immediate contour of the saccade endpoints (Δ1: 0.9776 > *ps* > 0.1782), and a similar spread of sensitivity across temporal delays for the other distances from the saccade endpoint. Also, as for the main analysis relative to the cue location, we found a significant improvement of sensitivity for discrimination targets shown between ~3.3° and ~4.7° from the saccade endpoint (Δ2) when comparing trials where the cue remained on the screen to trials where the cue offset preceded the saccade by approximately 950 ms (t0: d’ = 1.09 ± 0.13 vs. t950: d’ = 1.35 ± 0.14, *p* <0.0044), but not if this delay was shorter (i.e. t350 and t650: 0.9776 > *ps* > 0.9410). Moreover, we found an earlier spread of attention for discrimination targets shown at the greatest tested distances from the saccade endpoint (Δ3), with significant differences found for trials in which the cue disappeared approximately 350 ms (t0: d’ = 0.92 ± 0.12 vs. t350: d’ = 1.12 ± 0.13, *p* <0.0170) and after longer delays (i.e. t650 and t950: both 0.0080 > *ps* > 0.0001). This illustrates, a clear dissociation with the spread of saccadic endpoints not mirroring the spread of attention.

**Fig. 4.**
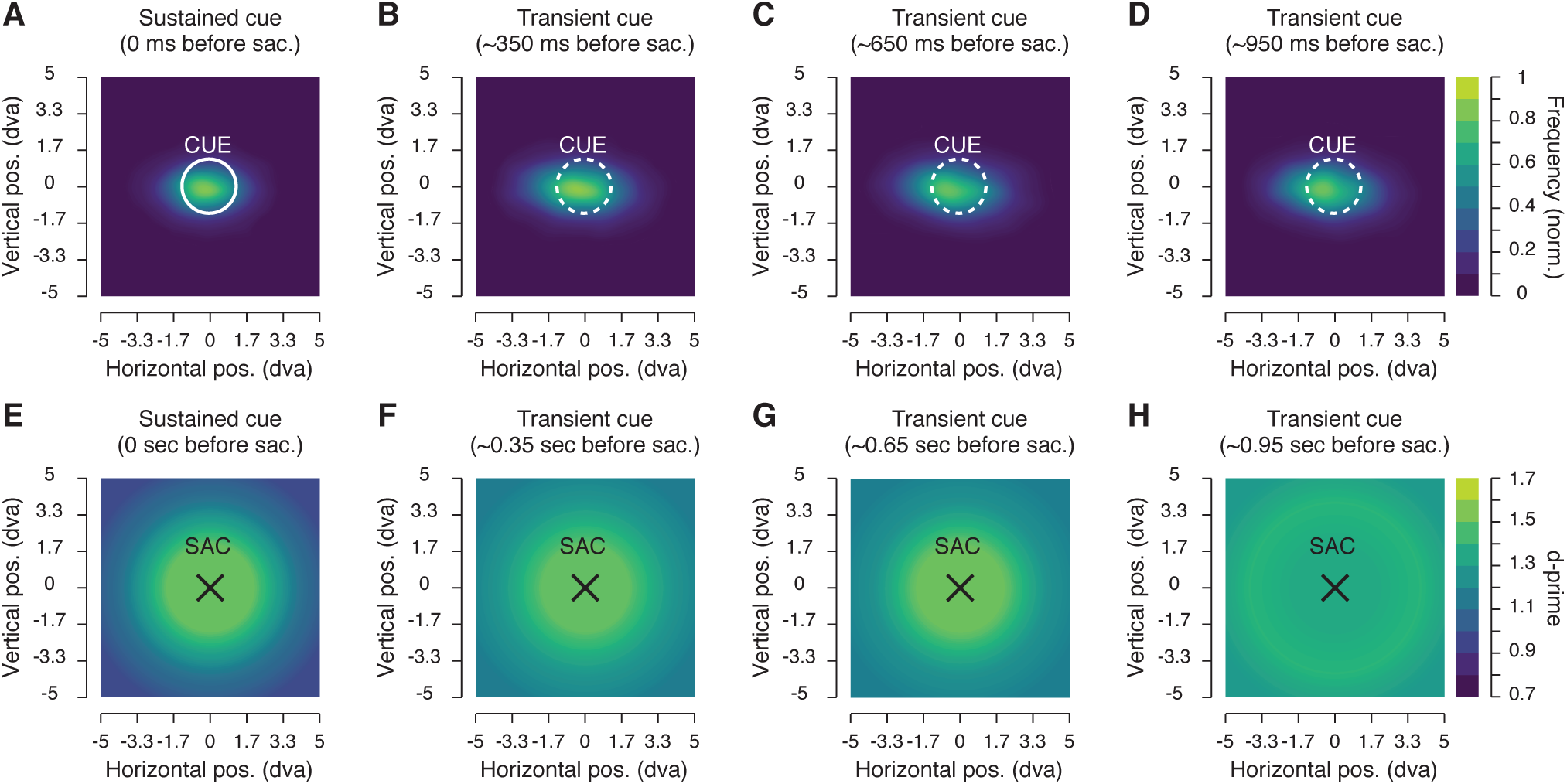
Saccade 3endpoint maps and sensitivity maps relative to saccade endpoint. **A-D**. Normalized saccade landing frequency maps averaged across participants. Data are shown for the sustained cue (A) and the transient cue conditions where the cue offset preceded the saccade by approximately 350 ms (B), 650 ms (C) or 950 ms (D). **E-H.** Each graph shows average sensitivity grouped in 3 distance (Δ1, Δ2 and Δ3) surrounding each trial saccade endpoint (SAC). Data are shown for the sustained cue (E) and the transient cue conditions for which the cue offset precede the saccade by approximately 350 ms (F), 650 ms (G) or 950 ms (H). Conventions are as in Figure 2.

## Discussion

We observed that when attention was allocated before an eye movement to a continuously present visual cue attentional resources remained bound to the cue location and its immediate surrounds (Δ1). Once the cue disappeared, we observed a spread of attention to more peripheral locations further away from the memorized target. This spread followed specific temporal dynamics. Notably, we found a modest although significant improvement in sensitivity at the farthest tested distance from the cue (Δ3) starting early after its offset. At a closer distance from the cue (Δ2), we observed a strong improvement in sensitivity beginning significantly later, about 650 ms after the cue offset. Interestingly, the spread of attention was never accompanied by a significant reduction in sensitivity at the most central tested locations, close to the cue location (Δ1). Here performance was always found to be highest, irrespective of whether the cue remained visible or disappeared from the screen. In other words, we did not observe a trade-off of attentional resources, where an increased spread of attention lead to a dilution of attention resources. Instead, we found that sensitivity increased overall (across all tested position) as the delay between the saccade and the cue offset grew. Furthermore, although saccades were more accurate when the cue remained on the screen, the increased spread of attention could not be explained by a spread of saccadic eye movements towards the target location. Although saccade accuracy slightly decreased when saccades were executed toward a cue that was no longer present, the sensitivity maps remained the same even after we corrected the tested position by the trial-by-trial retinal error. This meant that changing the spatial reference of the sensitivity maps from the center of the cue to the actual saccade endpoint made no difference, saccade landing then having no influence on the spread of attention. Our findings suggest that the deployment of attention before eye movements depends on the presence of the visual cue, with a concentration of attentional resources confined within ~2.4° when the cue remained on the screen and an undifferentiated spread of attention within ~4.7° around a cue that has disappeared about a second before. Through the assessment of sensitivity maps, we were able to capture a yet undocumented mechanism at play when attention no longer relies on the presence of a visual cue: the spread of attentional resources without trade-off.

The extent to which an exogenous cue modulates the deployment of spatial attention has been investigated previously, with outcomes mostly reflecting the difficulty of the task (Intriligator & Cavanagh, 2001). Our new paradigm allowed us to adjust the task difficulty to each participant and to control for visual eccentricity effects (Paradiso & Carney, 1988), two essential requirements for assessing visual sensitivity over space (Koenig-Robert & VanRullen, 2011). By combining a discrimination and a saccade task, we were able to verify through different saccade metrics that our measure of attention was not affected over space by the use of transient stimuli (Deubel, 2008; Deubel & Schneider, 1996). By analyzing the distribution of saccade endpoints we were able to demonstrate that participants effectively kept track of the cued position at all the tested delays.

We observed that the modulation of attention was tightly limited to the intended saccade target and its immediate surrounds when the target remained on the screen, with highest sensitivity observed at roughly an equivalent distance (10% of saccade size) to that observed in previous reports (Deubel & Schneider, 1996). Although performance was better than chance at more peripheral positions from the cue, suggesting that attention was not only allocated to the saccade target (Castet, Jeanjean, Montagnini, Laugier, & Masson, 2006; Kowler et al., 1995; Montagnini & Castet, 2007), we attributed this effect to our adjustment procedure and to the use of transient discrimination targets. Indeed, to adjust the difficulty of the task, we employed a threshold procedure to determine for each participant the specific discrimination target angles necessary for each eccentricity. Moreover, to evaluate the deployment of attention in an unstructured visual field we had to use transient targets, known to capture by themselves attention (Müller & Rabbitt, 1989; Theeuwes, 1991). With such a protocol, performance was thus maintained artificially high across space, allowing us to make conclusions only on the modulation of covert attention at different positions, rather than about absolute attention capacities and attentional deployment.

While our findings perfectly match those of studies which have investigated the allocation of attention with a continuously presented cue (Castet et al., 2006; Deubel, 2008; Deubel & Schneider, 1996; Kowler et al., 1995; Li, Barbot, & Carrasco, 2016;Montagnini & Castet, 2007; Rolfs & Carrasco, 2012; Rolfs, Jonikaitis, Deubel, & Cavanagh, 2011), to our knowledge, all studies investigating this aspect of attention used a structured visual field of visual placeholders. Despite using visual stimuli to measure attention we ensured these were presented only right before the saccades, at a time when the eye movements could no longer be stopped (Becker & Jürgens, 1979; Hanes & Schall, 1995) and when attention had already been allocated at the saccade target (Castet et al., 2006; Deubel, 2008; Klapetek, Jonikaitis, & Deubel, 2016; Li et al., 2016; Rolfs et al., 2011; Rolfs & Carrasco, 2012). We therefore captured for the first time the allocation of pre-saccadic attention in an unstructured visual field and observed a clear spread of attention from the cue center to its periphery. We then found that, contrary to the narrow allocation of presaccadic attention for a structured visual field, attention dynamically spread over space as a function of the delay between cue-offset and saccade onset. These effects are in contrast with a recent report in which the disappearance of a briefly presented cue, during a fixation task, lead to slower reaction times in detecting the presence of a flashed target on a black screen in the absence of placeholders (Taylor et al., 2015). Contrary to these effects, we observed an overall increase in sensitivity after the disappearance of the target. In our view, this difference is principally due to the difficulty in drawing conclusions about the deployment of attention based on the use of reaction times to supra-threshold stimuli (Handy et al., 1996). Indeed, with or without visual placeholders, reaction time benefits have been shown to cover entire visual quadrants or even entire visual hemifields (Bennett & Pratt, 2001; Hughes & Zimba, 1987; Taylor et al., 2015). These results are in contradiction with the tight allocation of attention to a saccade target, or to an exogenous cue, observed through changes in visual sensitivity (Deubel & Schneider, 1996; Handy et al., 1996). They are also in contradiction with findings showing changes in firing rate for attended stimuli placed inside, rather than outside visual and movement receptive fields (Gregoriou, Gotts, & Desimone, 2012; Moran & Desimone, 1985; Wurtz & Mohler, 1976).

Contrary to the notion of attention being a limited resource, we showed here that following the cue’s disappearance, attention spread without any trade-off between the cue’s center and its surround. We propose that this effect reflects receptive fields’ spatiotemporal dynamics; in particular, the transition from a modulated state, where receptive fields are directed toward the visible attended cue to a disengaged state and a return to their default position when the cue disappear. Across several studies, it was shown that attention modulates the response of visual cells’ receptive fields by shifting their spatial tuning toward a position closer to an attended target, whether this was a visible cue (Anton-Erxleben et al., 2009; Connor et al., 1997; Kay et al., 2015; Klein et al., 2014; Moran & Desimone, 1985; Niebergall et al., 2011; Sprague & Serences, 2013; Womelsdorf et al., 2006; 2008) or a visible saccade target (Neupane et al., 2016; Tolias et al., 2001; Zirnsak et al., 2014). But if receptive fields’ tuning can migrate toward a visible cue or respond to stimuli presented at locations outside their classical receptive field, what happens when the disappearance of the cue occurs? Once that happens, should the receptive fields not also be capable of returning to their resting-state position? Here, we hypothesize that following the cue’s offset, the receptive fields’ transition, back to their resting-state position, may result in a behavioral spread of attention toward more peripheral locations. It does not seem plausible that this dynamic change should occur instantaneously, and it may depend on the distance between the resting-state receptive field position and the attention field center. While electrophysiological evidence will be required to verify this hypothesis, the aforementioned interpretation embodies our beliefs for two principal reasons. First, a link was recently demonstrated between the cells’ receptive field spatial tuning and sensitivity employing a variety of attention tasks, for example, motion stimuli in animals (Niebergall et al., 2011) and face stimuli in humans (Kay et al., 2015; see also Sprague, Saproo, & Serences, 2015). The change in sensitivity measured over time in our task could thus reflect changes in neurons’ receptive field positions. Second, different studies have demonstrated strong connectivity between oculomotor and feature maps preceding the execution of a saccade (Gregoriou et al., 2012; Gregoriou, Gotts, Zhou, & Desimone, 2009; Moore & Armstrong, 2003). We propose that at the onset of the cue, both oculomotor cells (e.g. within the Frontal Eye Fields or the Superior Colliculus) and cells sensitive to visual features (e.g. within the primary visual cortex) shifted their tuning function toward the cue. At the cue offset, through sustained firing rate during the saccade delay (Gregoriou et al., 2012), oculomotor cells accurately maintain the spatial position of the cue. Just before the saccade, these oculomotor cells may send top-down signals to their retinotopically corresponding cells encoding visual features of the scene (Moore & Armstrong, 2003). Contrary to the oculomotor cells, these cells are principally driven by the presence of the cue. After the cue’s disappearance, they should therefore return to their resting-state position. Such transition between a modulated and a resting-state position may then drive our behavioral effect, where we observed an increase of sensitivity for discrimination targets shown at positions further away from the cue. We thus describe here a new mechanism of attention, which matches our experimental observations and predicts a spread of attention without an important trade-off of attentional resource between the attended location and its surround. Indeed, it would be expected that feature selective cells centered on the retinal position of the cue stay tuned to this attended position after the cue offset, maintaining high sensitivity within its vicinity. Moreover, as oculomotor cells in the current study were not driven by the visual signal but by the delayed oculomotor plan, there was no reason for saccade landing distribution to be correlated with the observed sensitivity spread.

We observed that the extinction of the cue results in an increase of the attentional field size, measured just before a saccade. Changes in attentional field size constitute the core of an influential computational model of attention (Reynolds & Heeger, 2009). This model proposes to reconcile contradictory empirical findings of the effects of attention on visual contrast sensitivity by a normalization process which combines a stimulus drive, a suppressive drive and an attention field. The model predicts a contrast gain (shift of the tuning contrast function) when the attention field size is large, relative to the attended stimulus, and a change in response gain (overall increase in response amplitude) when the attention field size is small, relative to the attended stimulus. As we did not test the effect of attention on contrast, but rather on orientation, and measured only performance for test stimuli at threshold, we cannot distinguish here between a shift of the orientation psychometric function and an overall increase in sensitivity. Nevertheless, we directly measured the extent of the attention field theorized in this model and observed a clear increase of the attention field size after a cue’s disappearance. Our reported effects moreover match with those reported in an fMRI study on delayed endogenous attention during fixation, in which Herrmann and colleagues (2010) found that attention field size increases when comparing trials with placeholders to those without, during a delay preceding the presentation of a cued target. Future studies should make use of such a fruitful manipulation to further test the predication of the normalization model of attention (Reynolds& Heeger, 2009). For example, using our behavioral methods of attention field size combined with systematic contrast change, one could observe a gradual change from a response gain to a contrast gain, as a function of the cue offset time relative to the saccade onset. This procedure could also be applied to other sensory domains and to multi-sensory integration, which is also thought to be processed through a similar normalization model (Carandini & Heeger, 2012).

Using a new paradigm, we evaluated the spatial and temporal dynamics of pre-saccadic attention following the disappearance of a cue. We found that pre-saccadic attention was mainly focused at the closest surround from the cue and spread to more peripheral locations as a function of the increasing delay between the cue offset and the saccade onset. This spread was accompanied by an overall increase in sensitivity over space that cannot be explained by a loss in spatial localization of the memorized location, as demonstrated through the assessment of saccade landing distributions. We therefore provided evidence here of a spread of attentional resources occurring without trade-off, that in our view can only be explained by visual receptive fields’ spatio-temporal dynamics following the cue disappearance.

## Method

### Participants

Twelve students of the LMU München participated in the experiment (age 19-29, 5 females, 11 right-eye dominant, 1 author), for a compensation of 10 Euros per hour of testing. All participants except one author (MP) were naive as to the purpose of the study and all had normal or corrected-to-normal vision. The experiments were undertaken with the understanding and written consent of all participants, and were carried out in accordance with the Declaration of Helsinki. The experiments were designed according to the ethical requirement specified by the LMU München and ethics approval for experiments involving eye tracking by the institutional review board.

### Setup

Participants sat in a quiet and dimly illuminated room, with their head positioned on a chin and fore-head rest. The experiment was controlled by an Apple Mac mini computer (Cupertino, CA, USA). Manual responses were recorded via a standard keyboard. The dominant eye’s gaze position was recorded and available online using an EyeLink 1000 Tower Mount (SR Research, Osgoode, ON, Canada) at a sampling rate of 1 kHz. The experimental software controlling the display, the response collection as well as the eye tracking was implemented in Matlab (The MathWorks, Natick, MA, USA), using the Psychophysics (Brainard, 1997; Pelli, 1997)} and EyeLink toolboxes (Cornelissen, Peters, & Palmer, 2002). Stimuli were presented at a viewing distance of 60 cm, on a 21-in gamma-linearized LaCie Electron 21/108 CRT screen (Paris, France) with a spatial resolution of 1,024 × 768 pixels and a vertical refresh rate of 120 Hz.

### Procedure

The study was composed of a main saccade task tested in 3 to 4 experimental sessions (on different days) of about 90 minutes each (including breaks). The main task was always preceded by a threshold task at the beginning of each experimental session. Each session was composed of 2 blocks of the threshold task followed by 3 to 4 blocks of the main saccade task. All participants ran a total of 20 blocks of the main saccade task.

### Main saccade task

Each trial began with participants fixating a central fixation target forming a black (~0 cd/m^2^) and white (88 cd/m^2^) “bull’s eye” (0.4° radius) on a gray background (44 cd/m^2^). When the participant’s gaze was detected within a 3.0° radius of a virtual circle centered on the fixation target, for at least 200 ms, the trial began with a fixation period of 500 ms. After this period, a cue consisting of a black (~0 cd/m^2^) outlined circle (1.25° radius, 0.1° width) was presented 10° to the right or to the left of the fixation target (see Figure 1). The cue either stayed on the screen for a duration of 500 ms (3/4 of the trials) or remained continuously on the screen until the end of the trial (1/4 of the trials). Participants were instructed to move their eyes as quickly and as accurately as possible toward the center of the cue at the offset of the fixation target, which occurred at different times after the cue onset (see below). Following the fixation target offset, one discrimination target and five distractors were shown for a duration of 25 ms. The positions of target and distractors were randomly selected among 25 possible positions homogeneously covering a 10° by 10° map centered on the cue (positions located at every second intersection of a 7 columns by 7 rows grid, see Figure 1C). All targets were Gabor patches (frequency: 1.75 cycles per degree; 100% contrast; random phase across trials; Gaussian envelope: 0.6°). While the distractors were vertical Gabors, the discrimination target was a tilted Gabor (clockwise or counter-clockwise relative to the vertical) with an angle adjusted in the threshold task for different distances from the fixation target (see threshold task). All targets were later replaced by Gaussian pixel noise masks for a duration of 25 ms (made of ~0.11°-width pixels with the same Gaussian envelope as the Gabors). We didn’t present any target or mask in 4% of all the trials, in order to evaluate the influence of our stimuli on the saccade execution (note that all other analyses are based on the discrimination target present trials).

At the end of each trial, participants reported the direction of the discrimination target using the keyboard (right or left arrow key) followed by a negative-feedback sound in the case of an incorrect response. On trials where no target was shown, participants randomly pressed one of the two response buttons, followed by a random feedback sound.

Participants completed between 2914 and 3773 trials of the main saccade task. Correct fixation resulted from gaze being maintained within a 3.0° radius virtual circle centered on the fixation target. Correct saccades resulted from saccades landing within a 4.0° radius virtual circle centered on the cue. Both criteria were checked online. Trials with fixation breaks or incorrect saccades were repeated at the end of each block, together with trials during which a saccade was initiated (crossing the virtual circle around the fixation target) within the first 50 ms or ended (crossing the virtual circle around the cue) after more than 350 ms following the fixation target offset (participants repeated between 114 to 973 trials across all sessions).

### Stimuli timing

The saccade signal delay (fixation target offset relative to the cue onset) was selected in order to have eye movement onset randomly interspersed between 700 and 1600 ms after the cue onset. In order to obtain this temporal range we had to account for systematic changes in saccade latencies, in function of the saccade signal delay. Indeed, in a pilot experiment we observed that participants quickly learned the range of possible delays, with shorter saccade latencies observed for longer signal delays and conversely. At the end of each block, we determined the slope (-0.09 ± 0.01 ms) and intercept (215.84 ± 4.88 ms) of a linear regression best describing this relationship (for the first block, we used fixed slope and intercept values of −0.5 ms and 200 ms, respectively). The saccade signal was then determined on each trial by subtracting the saccade latency estimated for each trial delay from a randomly selected duration (between 700 and 1600 ms in steps of ~8.3 ms, a screen frame). This gave us signal delays ranging between 428.92 ± 6.46 ms and 1435.33 ± 2.46 ms after the cue onset (from the trials after data pre-processing, see below).

Next, in order to probe attention in the last 100 ms preceding saccades we played the discrimination and distractors randomly between 50 and 100 ms (in steps of 25 ms), before the saccade latency, estimated for a given saccade signal delay trial. This resulted in discrimination target offset time relative to the saccade onset of **-**48.02 ± 1.16 ms (from the trials after data pre-processing, see below).

### Threshold task

The threshold task preceded the saccade main task at the beginning of each experimental session. This task made it possible to counteract possible learning effects and to adjust the baseline performance, for the presentation of a discrimination target at different distances from fixation, across participants. This latter point was particularly important as it reduced the impact of eccentricity effects (Paradiso & Carney, 1988) onto the mapping of attention benefits.

Contrary to the main task, participants were instructed to keep fixation on the fixation target, which remained on the screen. Also, compared to the main task, the cue could be presented at any of the 25 positions where the discrimination targets were shown, and it remained on the screen until the end of each trials. The discrimination target always followed the cue onset by 200 ms and was always presented at the cued location. With the exception of these differences the threshold task otherwise matched the main task.

The 25 possible positions of the discrimination target and cue were subdivided into 4 equiprobable groups of distances from the fixation target (distance 1: from ~5.3° to ~7.5°; distance 2: from ~8.5**°** to ~10.5°, distance 3: from ~11.8° to ~13.7°; distance 4: from ~15.1° to ~15.8°). Following a procedure of constant stimuli, the orientation of the discrimination target varied randomly across trials between five linearly spaced steps (between ±1° and ±17° for distances 1-2; and between ±1° and ±29° for distances 3-4).

Participants were instructed that the cue would always indicate the discrimination target’s location in all the trials and were told to report its orientation (clockwise or counter-clockwise) at the end of each trial. They completed 2 blocks of 160 trials, and correct fixation within a 3.0° radius virtual circle centered on the fixation target was checked online. Trials with fixation breaks were immediately discarded and repeated at the end of each block.

For the four main distances from the fixation target, we individually determined, for each participant and on each experimental session, four threshold values, corresponding to the discrimination target’s angles that would lead to correct discrimination on 85% of trials. To do so, we fitted four cumulative Gaussian functions to performance gathered in the threshold blocks. These threshold angles were used in the main task for discrimination targets, played at their respective distances from the fixation target.

### Data pre-processing

Before proceeding to the analysis of the behavioral results of the main task we scanned the recorded eye-position data offline. Saccades were detected based on their velocity distribution (Engbert & Mergenthaler, 2006) using a moving average over twenty subsequent eye position samples. Saccade onset and offset were detected when the velocity exceeded and fell behind the median of the moving average by 3 SDs for at least 20 ms. We included trials where a correct fixation was maintained within an 3.0° radius centered on the fixation target, where a correct saccade started at the fixation target and landed within an 4.0° radius centered on the cue and where no blink occurred during the trial. Finally, only trials in which the discrimination target offset occurred in the last 150 ms preceding saccades were included in the analysis. In total, we included 27414 trials (81.25% of the online selected trials, 69.48% of all trials played) of the main saccade task.

### Behavioral data analysis

We designed our experiments in order to have a similar amount of trials between 4 main delays, defined by the offset time of the cue relative to the onset of the saccade (t). We considered the condition in which the cue remained continuously on the screen as our first delay (t0) and next determined three bins of trials; one when the cue offset occurred between 200 and 500 ms (t350), one between 500 and 800 ms (t650) and one between 800 and 1100 ms (t950) before the saccade onset. For each participant and each of these conditions, we determined the sensitivity in discriminating the orientation of the discrimination target (d’):*d’* = *z*(hit rate) - *z*(false alarm rate). To do so, we defined a clockwise response to a clockwise discrimination target (arbitrarily) as a hit and a clockwise response to a counter-clockwise discrimination target as a false alarm. Corrected performance of 99% and 1% were substituted if the observed proportion correct was equal to 100% or 0%, respectively. Performance values below the chance level (50% or d’ = 0) were transformed to negative d’ values. Sensitivity was computed either separately for each of the 25 different position of the discrimination targets or for target positions grouped in three main distances (Δ) from the cue location (Δ1: from 0° to ~2.4°; Δ2: from ~3.3° to ~4.7°, Δ3: from ~5.3° to ~7.1°). These distances were arbitrarily defined in order to increase the power of our analyses. In our analyses across participants we included 40.76 ± 1.26 trials per discrimination target position and time condition, giving then 176.65 ± 5.47 trials per time condition when individual positions were combined in three main distances. We also computed sensitivity for test positions relative to the saccade landing position. To do so, for each trial we recomputed the distance between the observed saccade landing and the discrimination target coordinates individually. Later, we grouped trials into the same 3 distances as above (Δ), but this time from the saccade landing point (Δ1: from 0° to ~2.4°; Δ2: from ~3.3° to ~4.7°, Δ3: from ~5.3° to ~7.1°).

Individual sensitivity maps of target discrimination (see Figure 2A-D, middle panels) were first obtained by interpolating (triangulation-based natural neighbor interpolation) the missing values located every two intersections of the 7 columns by 7 rows grid. This was achieved by using the mean sensitivity for each participant, obtained over 25 positions of the discrimination target. Then the grid was rescaled (Lanczos resampling method) so as to obtain a finer spatial grain. These maps where then produced by drawing colored squares centered on their respective coordinates and following a linear color scale going from d’= 0.7 to d’=1.7. Groups of discrimination target sensitivity maps (see Figure 3A-D bottom panels and Figure 4E-H) were obtained by interpolating (linear interpolation) the mean sensitivity obtained over in the 3 different groups of main distances between the discrimination targets and the cue positions (Δ) or between the discrimination targets and the saccade endpoint position (Δ). These maps were then produced by drawing colored circles centered on the cue or saccade landing point, with a radius corresponding to their respective distances from the cue or the saccade landing point and following the same color scale used for the position sensitivity maps. A similar procedure was used to draw the threshold angle map (see Figure 1D) this time with a linear color scale going from threshold angles of 0° to 25° and colored circles centered on the fixation target. Saccade endpoint maps (Figure 4A-D) were obtained through the use of a bivariate kernel density estimator (Botev, Grotowski, & Kroese, 2010) were first normalized relative to the total amount of trials within a condition and later averaged across participants.

For statistical comparisons we drew 10,000 bootstrap samples (with replacement) from the original pair of compared values. We then calculated the difference of these bootstrapped samples and derived two-tailed *p* values from the distribution of these differences.

## Author contributions

Conceptualization, M.P., H.D. and M.S.; Methodology, M.P., H.D. and M.S.; Software, M.S.; Formal analysis, M.S.; Investigation, M.P.; Resources, M.S.; Writing – Original Draft, M.P. and M.S.; Writing – Review & Editing, M.P., H.D. and M.S.; Visualization, M.S.; Supervision, M.S.; Project administration, M.S.; Funding Acquisition: M.P. and M.S.

## Acknowledgments

The authors declare no competing financial interests. This research was supported by the Swiss National Foundation (MP: 100014_140379), a travel grant from “Fondation Ernst & Lucie Schmidheiny” in Geneva to M.P. and a “Deutsche Forschungsgemeinschaft” temporary position for principal investigator grant to M.S. (SZ343/1). We are grateful to the members of the Deubel laboratory in Munich for helpful comments and discussions and to Elodie Parison, Alice and Clémence Szinte for their invaluable support.

